# Quantitative Three-dimensional Label-free Digital Holographic Imaging of Cardiomyocyte Size, Ploidy, and Cell Division

**DOI:** 10.1101/2023.11.02.565407

**Authors:** Sangsoon Park, Herman Huang, Ines Ross, Joseph Moreno, Sheamin Khyeam, Jacquelyn Simmons, Guo N. Huang, Alexander Y. Payumo

**Affiliations:** Cardiovascular Research Institute & Department of Physiology, University of California, San Francisco, San Francisco, CA, 94158, USA; Eli and Edythe Broad Center for Regeneration Medicine and Stem Cell Research, University of California, San Francisco, San Francisco, CA, 94158, USA; BAKAR Aging Research Institute, University of California, San Francisco, San Francisco, CA, 94158, USA; Department of Biological Sciences, San Jose State University, San Jose, CA 95192, USA

## Abstract

Cardiac regeneration in newborn rodents depends on the ability of pre-existing cardiomyocytes to proliferate and divide. This capacity is lost within the first week of postnatal development when these cells rapidly switch from hyperplasia to hypertrophy, withdraw from the cell cycle, become binucleated, and increase in size. How these dynamic changes in size and ploidy impact cardiomyocyte proliferative potential is not well understood. In this study, we innovate the application of a commercially available digital holographic imaging microscope, the Holomonitor M4, to evaluate the proliferative responses of mononucleated diploid and binucleated tetraploid cardiomyocytes. This instrument coupled with the powerful Holomonitor App Suite software enables long-term label-free quantitative three-dimensional tracking of primary cardiomyocyte dynamics in real-time with single-cell resolution. Our digital holographic imaging results provide direct evidence that mononucleated cardiomyocytes retain significant proliferative potential as most can successfully divide with high frequency. In contrast, binucleated cardiomyocytes exhibit a blunted response to a proliferative stimulus with the majority not attempting to divide at all. Nevertheless, some binucleated cardiomyocytes were capable of complete division, suggesting that these cells still do retain limited proliferative capacity. By quantitatively tracking cardiomyocyte volume dynamics during these proliferative responses, we reveal that both mononucleated and binucleated cells reach a unique size threshold prior to attempted cell division. The absolute threshold is increased by binucleation, which may limit the ability of binucleated cardiomyocytes to divide. By defining the interrelationship between cardiomyocyte size, ploidy, and cell cycle control, we will better understand the cellular mechanisms that drive the loss of mammalian cardiac regenerative capacity after birth.

## INTRODUCTION

Newborn rodents possess a transient capacity for cardiac regeneration that is lost shortly after birth (Porrello et al., 2011; H. Wang et al., 2020). Heart regeneration during this postnatal period relies on the ability of pre-existing cardiomyocytes to enter the cell cycle, divide, and replace damaged cardiac muscle tissue (Porrello et al., 2013). However, within this first week of life, rodent cardiomyocytes rapidly transition from hyperplasia to hypertrophy as they permanently withdraw from the cell cycle, undergo polyploidization, and increase in size (Bishop et al., 2021; Clubb & Bishop, 1984; Li et al., 1996; Soonpaa et al., 1996; Zak, 1973). During this transition, mononucleated diploid cardiomyocytes enter the cell cycle but fail to complete cytokinesis resulting in binucleated tetraploid cells (Engel et al., 2006; Leone et al., 2018; Leone & Engel, 2019; Stopp et al., 2017). How increased cell size and ploidy impacts cardiomyocyte cell cycle regulation and thus cardiac regenerative potential remain incompletely understood.

Cardiac primary cell culture represents a powerful *in vitro* approach to complement *in vivo* studies. Experiments using primary cardiomyocyte cultures from newborn rodents have revealed many intrinsic (e.g., transcription factors) and extrinsic (e.g., extracellular matrix proteins, secreted factors) regulators of cardiomyocyte proliferation and hypertrophy (Mehdipour et al., 2023). Therefore, primary cardiac cell culture is a relevant approach to investigate questions regarding heart development, physiology, and pathophysiology, especially at cellular and molecular levels.

A key challenge in studying cardiomyocyte cell cycle regulation is the difficulty to distinguish true cardiomyocyte cell division from cytokinesis failure and polyploidization both *in vitro* and *in vivo* (Auchampach et al., 2022). Traditional approaches rely on tissue fixation and staining for commonly used cell cycle markers (e.g., Ki67, PHH3, AurkB, etc.). However, the expression of these markers in cardiac muscle cells can often be misleading since they are similarly expressed in both cardiomyocytes that complete and fail cell division. Therefore, direct live-cell imaging of primary cardiomyocytes is currently the most definitive approach to demonstrate true cardiomyocyte division and expansion *in vitro* (Leone et al., 2018; Leone & Engel, 2019; Yahalom-Ronen et al., 2015).

In this study, we innovate the application of a commercially available digital holographic phase imaging microscope, the HoloMonitor M4 (Phase Holographic Imaging; Lund, Sweden), to investigate how size and ploidy influence the outcomes of cardiomyocyte cell division *in vitro*. The Holomonitor M4 is designed to fit within standard cell culture incubators and requires no additional environmental controllers for long-term imaging of living cells (**Fig. 1A**). The interference pattern created between a sample and reference beam generates phase shift data, which is then reconstructed to visualize the cultured cells in three dimensions (**Fig. 1B**). This system has several advantages compared to existing live-cell imaging technologies (Mölder et al., 2008). First, imaging is performed with a low-power 635 nm laser, which minimizes phototoxicity to allow gentle, label-free, long-term imaging of cellular behaviors. Second, full three-dimensional reconstructions are automatically generated for cells located within a 40 μm distance from the cell culture surface, which greatly ameliorates issues with maintaining sample focus during extended imaging durations. And third, the Holomonitor App Suite analysis software enables quantitative tracking of cell size, shape, and movement with single-cell resolution. These technological advantages empower us to quantitatively define cell volume dynamics that accompany successful and failed cell division in both mononucleated and binucleated cardiomyocytes. Using this approach, we explore the interrelationship between cardiomyocyte size, ploidy, and cell cycle control to better understand the cellular mechanisms that drive the loss of mammalian cardiac regenerative capacity after birth.

**Figure 1:**
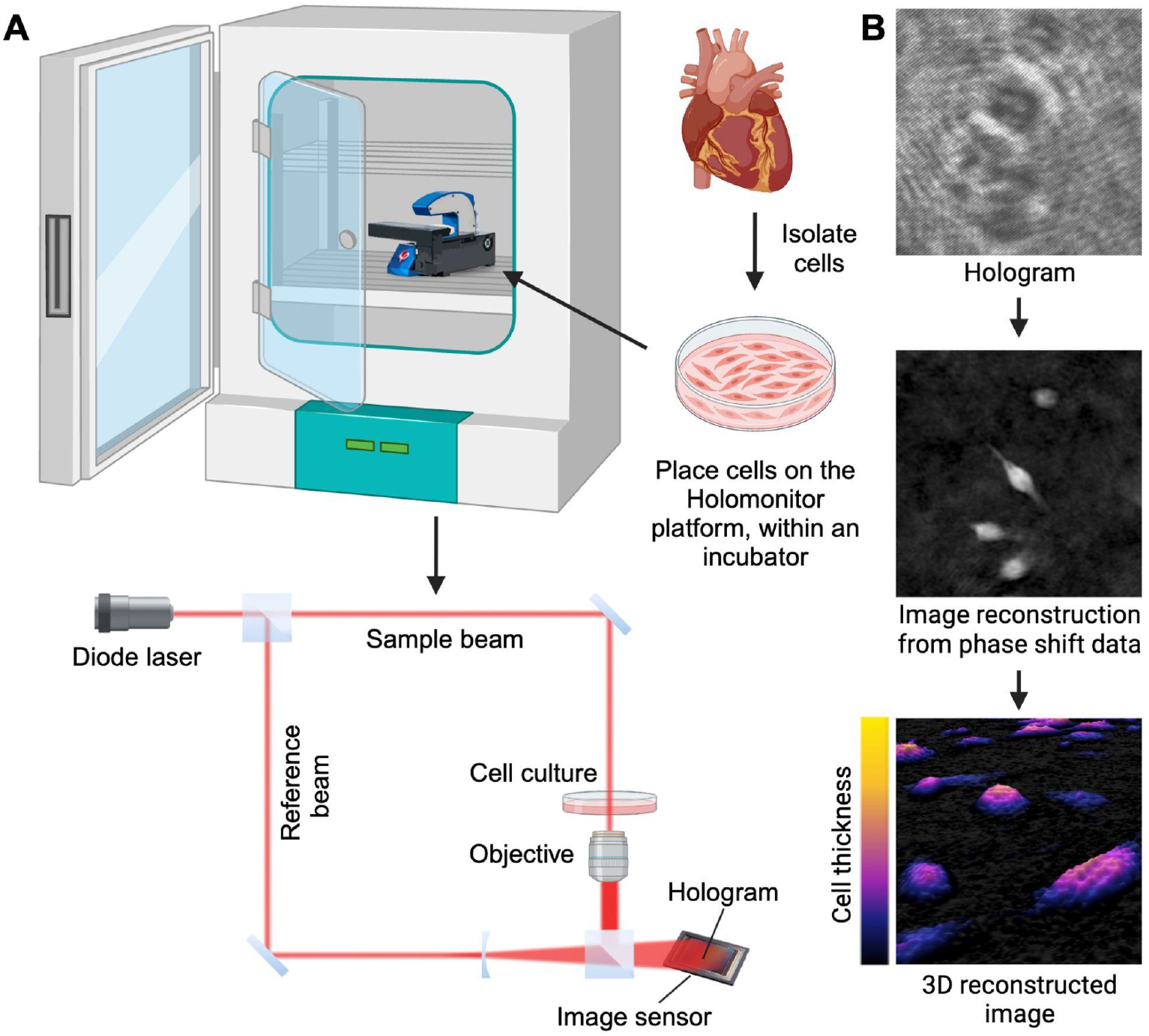
Three-dimensional digital holographic imaging of cultured primary cardiac cells with the Holomonitor M4. (**A**) Primary cardiac cultures are placed onto the Holomonitor M4 housed within a standard CO_2_ incubator. A diode laser is divided into a reference beam and an object beam that passes through the sample, which creates a phase shift. An interference pattern hologram is generated by merging of the object and reference beams. This is recorded by an image sensor to reconstruct a phase image. (**B**) Example images of an interference pattern hologram and three-dimensional reconstructions of cultured primary rat cardiomyocytes as visualized on the Holomonitor M4. Created with BioRender.com.

## RESULTS

### Primary cardiomyocytes are distinguished from non-cardiomyocytes based on their larger size and limited motility

Primary rodent cardiomyocyte cultures are often contaminated with non-cardiomyocytes including cardiac fibroblasts and endothelial cells. While methods to improve cardiomyocyte purity have been developed based on differential attachment speeds (Blondel et al., 1971) or densities (Sen et al., 1988), the addition of these enrichment steps nevertheless fail to completely remove all non-cardiomyocytes in the culture. Therefore, live-imaging experiments of primary cardiomyocyte cultures often necessitates direct cardiomyocyte-specific labeling or post-imaging fixation and validation of cardiomyocyte identity by immunostaining, which adds to experimental complexity (Leone & Engel, 2021). The HoloMonitor M4 enables direct quantitative three-dimensional label-free imaging of any adherent cells in culture. Cardiomyocytes make up most of the heart’s volume due to their larger size but are outnumbered by smaller cardiac fibroblasts and endothelial cells (Baum & Duffy, 2011). We therefore evaluated if our label-free digital holographic imaging approach could differentiate primary cardiomyocytes from non-cardiomyocytes in mixed primary cardiac cultures based on their unique three-dimensional morphologies and dynamics.

To define which primary cardiac cells observed on the Holomonitor M4 corresponded to cardiomyocytes, we first live-imaged the culture using digital holographic microscopy, fixed the cells, immunostained against the cardiomyocyte-specific marker α-actinin, and then re-imaged the same field by conventional fluorescence microscopy (**Fig. 2A-B**). The thicker and larger cells identified on the Holomonitor M4 most strongly correlated with α-actinin-positive cardiomyocytes (**Fig. 2B-C**). The maximum optical thickness (**Fig. 2D**) and optical volume (**Fig. 2E**) of these putative cardiomyocytes were 1.8-fold and 3.2-fold higher than non-cardiomyocytes, respectively. While size alone was a strong predictor of cardiomyocyte identity in most cases, we noticed a few thicker α-actinin-negative non-cardiomyocytes by digital holographic imaging (**Fig. 2A-B**). Because of this, we evaluated cellular motility as a secondary identification criterion and discovered that primary rat cardiomyocytes are slow moving, covering 6.7-fold less distance than non-cardiomyocytes within a 12-hour period (**Fig. 2F-H; Movie S1**). These data demonstrate that our label-free digital holographic imaging approach can reliably distinguish primary neonatal rat cardiomyocytes from non-cardiomyocytes based on their larger size and limited motility. We additionally found that these criteria can be similarly applied to identify primary mouse cardiomyocytes as well (**Fig. S1**), broadening the applicability of this approach.

**Figure 2:**
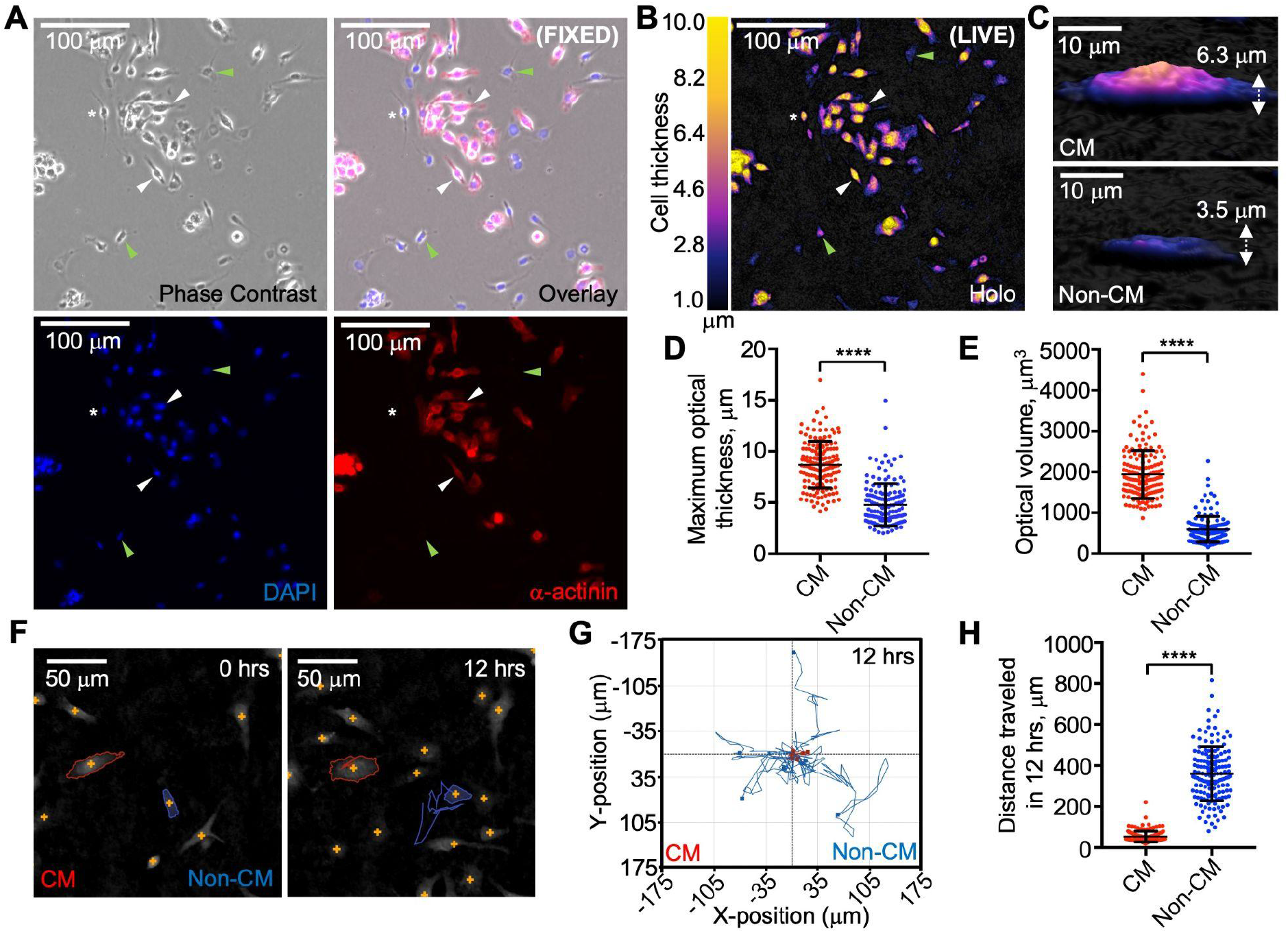
Label-free digital holographic microscopy identifies primary rat cardiomyocytes based on their larger size and limited motility. (**A-B**) The same imaging field of fixed postnatal day 2 (P2) rat cardiac cells visualized on a phase contrast fluorescence microscope (**A**) or live on the Holomonitor M4 (**B**). Cardiomyocytes (CM) are α-actinin-positive (**A**, white arrowheads) and larger (**B**, white arrowheads) while non-CMs are α-actinin-negative (**A**, green arrowheads) and smaller (**B**, green arrowheads). DAPI labels all nuclei. White asterisks highlight an α-actinin-negative non-CM that appears thicker. (**C-E**) Representative three-dimensional reconstructions of a CM and non-CM (**C**) and quantifications of their maximum optical thicknesses (**D**) and optical volumes (**E**). (**F-H**) Motility analysis of CMs and non-CMs. (**F)** Representative segmentation and tracking of a CM (red outline) and non-CM (blue outline) over 12 hours. Cell trajectories are shown as a single solid line for each cell. (**G**) Overlayed trajectories of CMs (n=7) and non-CMs (n=7) over 12 hours. All cell traces begin at the (0,0) coordinate. (**H**) Total distance traveled by each cell over 12 hours. 150 CMs and 151 non-CMs from postnatal day 2 (P2) hearts were analyzed in **D**, **E**, and **H**. Values are reported as mean ± standard deviation. Student’s t-test; ****, *P* < 0.0001.

### Digital holographic imaging permits quantitative tracking of cardiomyocyte volume dynamics with single-cell resolution

Cardiomyocytes undergo hypertrophy to maintain cardiovascular function in response to overload (Nakamura & Sadoshima, 2018), and this response can be modeled *in vitro*. For example, treatment of cultured primary rat cardiomyocytes with the sympathetic neurohormone norepinephrine results in hypertrophic growth through activation of α_1_-adrenergic receptors (Simpson, 1985). Phenylephrine is a selective α_1_-adrenergic receptor agonist that promotes cardiomyocyte hypertrophy by inducing protein synthesis via various pathways including MAPK/ERK (Rolfe et al., 2005) and mTOR signaling (Hannan et al., 2003). We thus evaluated the quantitative capacities of the HoloMonitor M4 to detect changes in three-dimensional cardiomyocyte volume after stimulation with phenylephrine.

Treatment with 50 µM phenylephrine for 48 hours increased the two-dimensional size of α-actinin-positive cardiomyocytes as visualized by conventional fluorescence imaging (**Fig. 3A-B**). We next employed our digital holographic imaging approach to quantify the three-dimensional optical volumes of individual cardiomyocytes prior to and after 48-hour treatment with phenylephrine (**Fig. 3C-F**). While no significant differences were observed in the optical volumes between untreated (n = 305) and treated (n = 300) cardiomyocytes at the start of the experiment (**Fig. 3D**), the average optical volume increased by 1.33-fold in treated cells after 48 hours (**Fig. 3E; Movie S2**). A powerful feature of the Holomonitor App Suite software is its image segmentation capabilities that enable three-dimensional tracking and analysis of single-cell dynamics (**Fig. 2F**). Single-cardiomyocyte tracking revealed heterogeneity in cardiomyocyte volume responses to the hypertrophic stimulus (**Fig. 3F**). For example, 48.2% of untreated cardiomyocytes decreased in volume over the 48-hour period in comparison to only 13% of cardiomyocytes stimulated with phenylephrine. 50.8% of untreated cardiomyocytes demonstrated a modest less than two-fold increase in optical volume while 75.3%, 9.3%, and 1.3% of phenylephrine treated cardiomyocytes exhibited a wider range in cell growth between 1- and 2-fold, 2- and 3-fold, and greater than 3-fold, respectively (**Fig. 3G**). These experiments illustrate a novel application of the Holomonitor M4 to dynamically track three-dimensional volume changes in individual primary cardiomyocytes over time, which provides deeper insights into the complexity of cardiomyocyte hypertrophic responses at the single-cell level.

**Figure 3:**
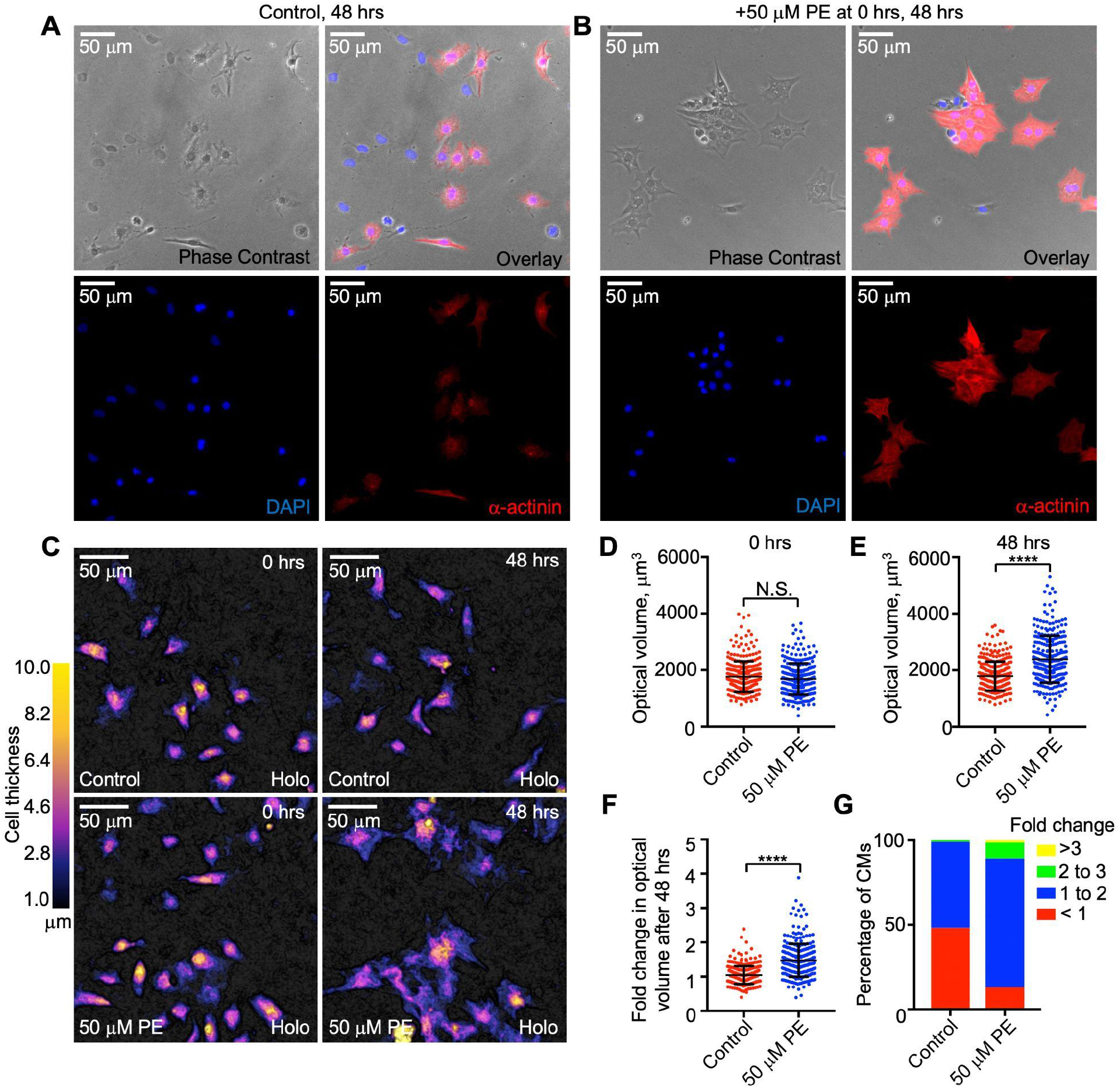
Digital holographic imaging of cardiomyocyte volume dynamics in response to a hypertrophic stimulus. (**A-B**) Phase-contrast and fluorescence micrographs of primary rat cardiomyocytes (CM) untreated (**A**) or treated with 50 µM phenylephrine (PE; **B**) for 48 hours. Cardiac cultures were stained for the CM-specific marker α-actinin and DAPI to label all nuclei. (**C-G**) Representative digital holographic images of untreated CMs or those treated with 50 µM PE for 48 hours (**C**). (**D-G**) Quantification of CM raw optical volumes at 0 hours (**D**) and 48 hours (**E**). Fold change in CM optical volumes observed after 48 hours (**F**). Percentage of CMs that exhibit fold changes in volume within specified bins (**G**). 305 untreated and 300 treated cardiomyocytes from postnatal day 1 (P1) and P2 hearts were analyzed. Values are reported as mean ± standard deviation. Student’s t-test; ****, P < 0.0001; N.S., not significant.

### Digital holographic imaging of cycling cardiomyocytes reveals both successful and failed attempts at cell division

In our base serum-free culture conditions, we evaluated the frequency of cardiomyocyte cell cycle entry by incubating primary neonatal rat cardiomyocytes in the thymidine analog 5-ethynyl-2’deoxyridine (EdU) for 48 hours, which identifies cells undergoing DNA synthesis (S-phase) within that time period. After immunostaining against the cardiomyocyte-specific marker cardiac troponin T (cTnT), we quantified the percentage of cardiomyocytes that incorporated EdU and determined that only 8.37% of these cells had entered S-phase (**Fig. 4A, 4C**). Because this low frequency of cardiomyocyte cell cycle entry would be challenging to monitor in our label-free digital holographic imaging experiments, we sought ways to better support robust cardiomyocyte proliferation *in vitro* while maintaining chemically-defined conditions.

**Figure 4:**
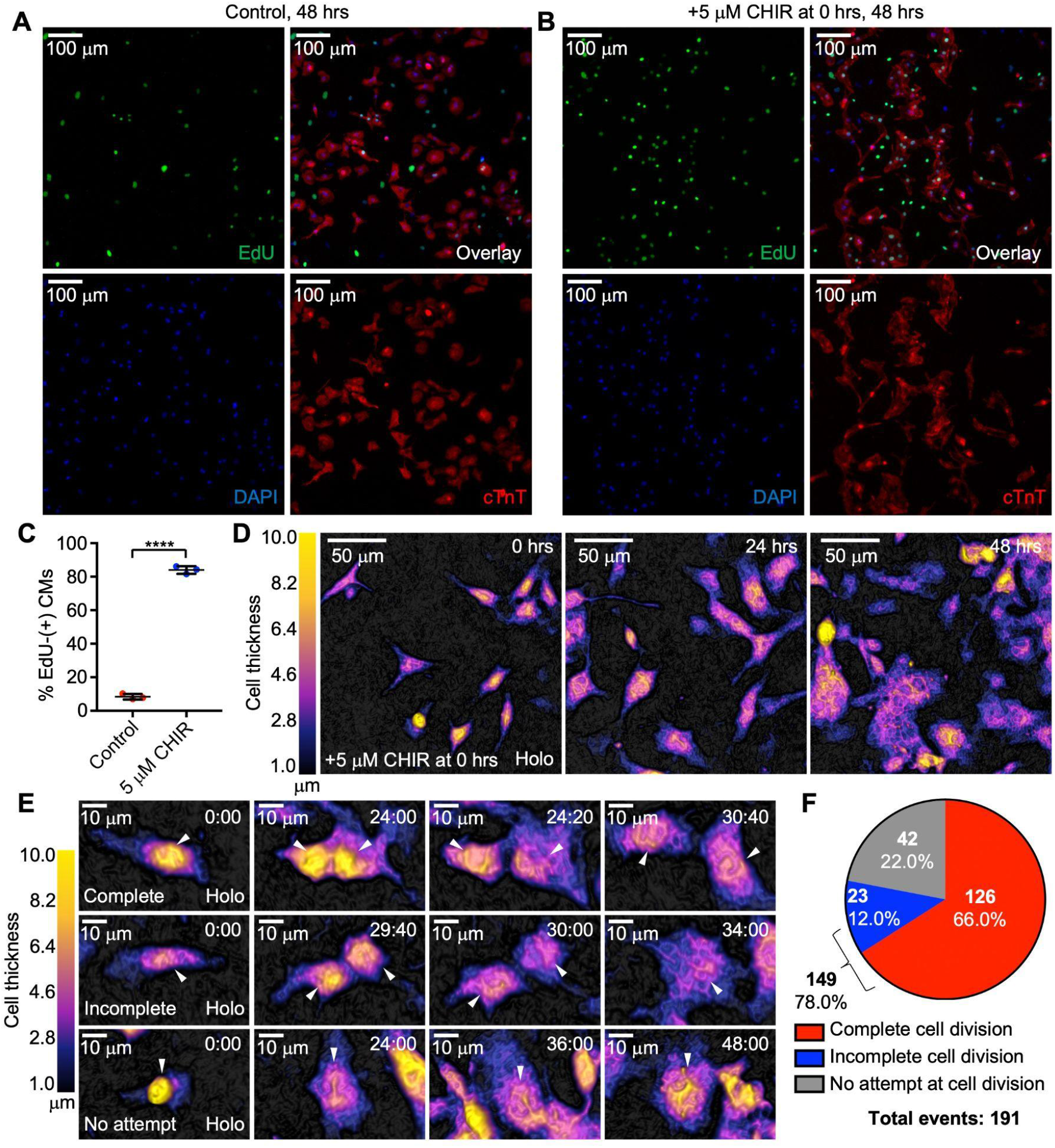
Digital holographic imaging distinguishes between complete and incomplete cardiomyocyte cell division. (**A-C**) Representative fluorescence micrographs of postnatal day 1 (P1) rat cardiomyocytes (CM) untreated (**A**) or treated with 5 µM CHIR99021, a GSK-3β inhibitor (**B**) for 48 hours. Cultures were stained for CM-specific marker cardiac Troponin T (cTnT) and DAPI to label all nuclei. Cells incorporating ethynyl-2’-deoxyuridine (EdU), an S-phase marker, after 48-hour incubation were identified. Quantification of EdU- and cTnT-double positive cells from three independent cultures (**C**). (**D**) Representative digital holographic images of P3 CMs stimulated with 5 µM CHIR99021 at 0 h, 24 h, and 48 hrs post-treatment. (**E-F**) Representative digital holographic time-lapse images showing examples of CMs that complete, incomplete, or make no attempts at cell division post stimulation (time stamp, hours:minutes) (**E**) and outcome analysis of 191 total CMs evaluated (**F**). Values are reported as mean ± standard deviation. Student’s t-test; ****, P < 0.0001.

Glycogen synthase kinase 3 (GSK3) has been proposed as a therapeutic target for cardiac regenerative medicine (Singh et al., 2019). Treatment of primary neonatal rat cardiomyocytes with the GSK3 inhibitor 6-bromoindirubin-30-oxime (BIO) resulted in a 10-fold increase in cardiomyocyte cell cycle entry (Tseng et al., 2006). More recently, the GSK3β-specific inhibitor CHIR99021 was discovered to increase the expression of cell cycle markers in primary cultured human cardiomyocytes through activation of Wnt/β-catenin signaling (S. Wang et al., 2016). Wnt signaling is required for cardiac progenitor expansion during embryonic development (Kwon et al., 2007), but may then be inactivated by Hippo signaling after birth to restrain cardiomyocyte proliferation and regulate heart size (Heallen et al., 2011). Therefore, we reasoned that activation of Wnt signaling through GSK3 inhibition would be a developmentally relevant stimulus to evaluate neonatal cardiomyocyte proliferative responses in our *in vitro* experiments.

Treatment of primary cardiac cultures with 5 µM CHIR99021 robustly increased cardiomyocyte-specific EdU incorporation to 84.0%, a 10-fold increase above that observed using our base media alone (**Fig. 4B, 4C**). During these 48 hours of continuous stimulation with CHIR99021, digital holographic imaging revealed dramatic changes in cardiomyocyte morphology and behavior (**Fig. 4D; Movie S3**), including notable increases in cardiomyocyte size. We quantified the single-cell behaviors of 191 randomly chosen cardiomyocytes to further understand their individual responses to the defined proliferative stimulus. We observed cardiomyocytes that 1) completed cell division, 2) failed cell division, or 3) made no attempt at cell division within the 48-hour experimental window (**Fig. 4E**). We examined the frequencies of these major outcomes and discovered that 66% of cardiomyocytes analyzed displayed complete cell division while 12% failed to completely divide, which suggests that 78% of all cardiomyocytes analyzed had entered the cell cycle. This overall frequency of cardiomyocyte cell cycle entry after treatment with CHIR99021 is in agreement with that predicted by our fixed-cell cardiomyocyte EdU incorporation experiments (**Fig. 4C**).

### Binucleated cardiomyocytes exhibit a blunted proliferative response but retain capacity for successful cell division

We noticed that it was possible to distinguish mononucleated from binucleated cardiomyocytes by identifying their nuclei based on their differential densities when compared to that of the surrounding perinuclear regions (**Fig. 5A**). During our digital holographic imaging experiments, cardiomyocyte nuclei could be identified as round structures with approximately 10 μm diameters that have less overall vertical thickness. Polyploidization is thought to limit the proliferative potential of cardiomyocytes and is considered a major barrier to cardiac regeneration (Derks & Bergmann, 2020; Huang et al., 2023; Mehdipour et al., 2023). Therefore, we next determined if mononucleated and binucleated cardiomyocytes exhibited different proliferative responses to CHIR99021 stimulation.

**Figure 5:**
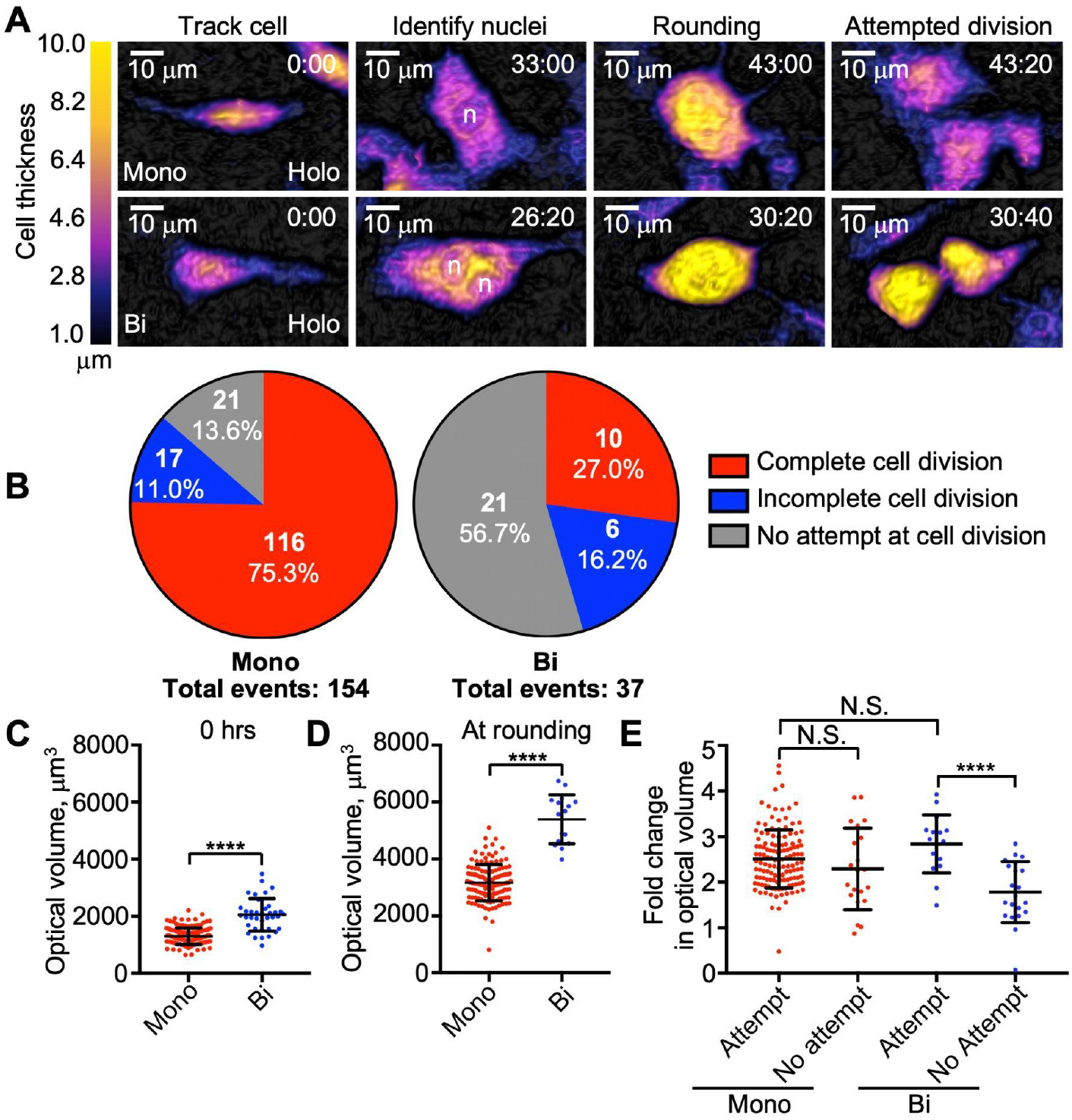
Binucleation blunts cardiomyocyte responses to a proliferative stimulus. (**A**) Representative digital holographic time-lapse images of postnatal day 3 (P3) cardiomyocytes (CMs) stimulated to divide with 5 µM CHIR99021 (timestamp, hours:minutes). Example nuclei identified to distinguish mononucleated from binucleated CMs are marked with “n”. (**B**) Outcome analyses of all 154 mononucleated and 37 binucleated CMs analyzed. (**C-D**) Optical volumes of all mononucleated and binucleated CMs in **B** at 0 hours (**C**), and the time of rounding (n = 133 and n = 16, respectively) (**D**). Fold change in optical volumes observed at rounding for mononucleated and binucleated CMs that attempt to divide (n = 133 and n = 16, respectively) or by 48 hours for mononucleated and binucleated CMs that make no attempt at cell division (n = 21 and n = 21, respectively) (**E**). Values are reported as mean ± standard deviation. (**C-D**) Student’s t-test; ****, P < 0.0001. (**E**) One-way ANOVA, Tukey’s multiple comparisons test; ****, P < 0.0001; N.S., not significant.

We subcategorized the 191 cardiomyocytes analyzed previously (**Fig. 4F**) into mononucleated (n = 151, 80.6%) and binucleated (n = 37, 19.4%) cardiomyocytes. These primary cardiac cultures were derived from postnatal day 3 (P3) neonatal rats, a developmental stage when cardiomyocytes are just beginning to binucleate *in vivo* (Li et al., 1996). Since our cultures contain a mixture of both mononucleated and recently binucleated cardiomyocytes derived from the same hearts, we are able to more directly resolve the unique behaviors of these cells under identical culture conditions without any further experimental manipulation. In response to 5 µM CHIR99021 treatment for 48 hours, we observed subpopulations of both mononucleated and binucleated cardiomyocytes attempting to divide. Strikingly, all observed events began with the cardiomyocyte rounding up and increasing in thickness within 20 minutes of attempted division (**Fig. 5A**), regardless of nuclei number. Mitotic rounding enabled by compliant extracellular matrices has been positively associated with complete cardiomyocyte cell division (Yahalom-Ronen et al., 2015). However, a recent report suggests that rounding up is not a marker to discriminate between complete cardiomyocyte division and binucleation (Leone et al., 2018). Our findings are consistent with the latter study and support that mitotic rounding itself is not a sufficient marker to predict true cardiomyocyte cell division.

Despite our observation that all cardiomyocytes analyzed rounded up prior to attempted cell division, not all cells were able to divide successfully. 86.3% of all mononucleated cardiomyocytes analyzed attempted to divide, with the majority, 75.3%, successfully completing cell division and 11.0% failing to completely separate (**Fig. 5B; Movie S4**). This finding supports the current hypothesis that mononucleated cardiomyocytes retain high proliferative and regenerative potential (Becker & Hesse, 2020). In contrast, only 43.2% of all binucleated cardiomyocytes analyzed attempted to divide with 27% successfully completing cell division and 16.2% failing cytokinesis (**Fig. 4B; Movie S5**). These results provide evidence that binucleation is indeed a barrier to cardiomyocyte proliferation. Our observations also demonstrate, however, that binucleated cardiomyocytes do at least retain some potential to successfully divide. This is in agreement with previous work documenting the successful division of binucleated rat cardiomyocytes *in vitro*, which may occur through the formation of pseudo-bipolar mitotic spindles (D’Uva et al., 2015; Leone & Engel, 2019; W. E. Wang et al., 2017).

### Cardiomyocyte binucleation increases a size threshold that may be required for attempted division

While only 13.6% of mononucleated cardiomyocytes did not attempt cell division in response to 5 µM CHIR99021 stimulation, the majority (56.7%) of binucleated cells analyzed made no attempt to do so (**Fig. 5B**). Previous studies have proposed that mammalian cells may need to meet a critical size threshold in order to exit from the G1 phase, progress through the cell cycle, and divide (Ginzberg et al., 2015). Increased ploidy is generally associated with increased cell size (Amodeo & Skotheim, 2016), and binucleated cardiomyocytes are noticeably larger than mononucleated cardiomyocytes (Windmueller et al., 2020). We wondered if cardiomyocytes that did not attempt to divide in response to CHIR99021 treatment failed to reach a critical size threshold to progress through the cell cycle. Therefore, we determined if such a size threshold exists in primary cardiomyocytes prior to attempted cell division and how binucleation influences this parameter.

We applied our digital holographic imaging approach to assess cell volume changes during the proliferative responses of both mononucleated and binucleated cardiomyocytes. We measured the initial optical volumes of P3 mononucleated and binucleated cardiomyocytes at the time of CHIR99021 stimulation. The average optical volumes of mononucleated and binucleated cardiomyocytes were 1304 +/− 279 μm^3^ and 2047 +/− 575 μm^3^, respectively (**Fig. 5C**), confirming that binucleated cardiomyocytes are initially larger than mononucleated cardiomyocytes on average. To determine the size of these same cells just prior to cell division, we measured cardiomyocyte optical volumes at the time of rounding (**Fig. 5D**). The average optical volumes of mononucleated and binucleated cardiomyocytes at rounding were 3122 +/− 704 μm^3^ and 5398 +/− 859 μm^3^, respectively. However, when evaluating the overall ratios by which these cells increased their volumes prior to attempted cell division, we noted that mononucleated and binucleated cardiomyocytes similarly enlarged by 2.47 +/− 0.67-fold and 2.84 +/− 0.64-fold, respectively (**Fig. 5E**). Taken together, these results show that binucleation not only increases initial cardiomyocyte size but also increases an absolute volume threshold reached prior to attempted cell division. Interestingly, the relative increase in cell volume is proportional in both mononucleated and binucleated cells.

We then further analyzed the individual responses of cardiomyocytes that made no attempt to divide despite stimulation with CHIR99021. The 21 mononucleated cardiomyocytes that did not attempt cell division within 48 hours still grew by 2.29 +/− 0.90-fold on average by the end of our video recordings, a similar volume increase exhibited by mononucleated and binucleated cells that had attempted division (**Fig. 5E**) within this time frame. This may suggest that the rate of cardiomyocyte growth may also play a role in determining if the cell will divide or that reaching a size threshold alone is not sufficient to trigger cell division. However, the 21 binucleated cardiomyocytes that did not divide within 48 hours demonstrated a statistically significant deficiency in growth, as their optical volumes only increased by 1.78 +/− 0.67-fold on average by the end of our videos (**Fig. 5E**). These findings suggest that binucleation delays cardiomyocyte growth in response to CHIR99021 treatment, which may result in their blunted proliferative response and lack of cell division.

### Initial cardiomyocyte size is not correlated with the timing of attempted cell division

We evaluated the timing of cardiomyocyte attempted cell division after stimulation with 5 μM CHIR99021 (**Fig. S2A**). Mononucleated cardiomyocytes that successfully or failed to divide attempted to do so within 1867 +/− 380 and 1805 +/− 386 minutes, respectively. Binucleated cardiomyocytes that successfully or failed to divide attempted to do so within 2170 +/− 464 and 2033 +/− 605 minutes, respectively. No statistically significant differences were observed in the timing of attempted cell division across all these conditions. Since binucleated and mononucleated cardiomyocytes exhibit proportional increases in volume prior to attempted division (**Fig. 5E**), this result suggests that binucleated cardiomyocytes must increase their volumes at a faster rate in response to CHIR99021 treatment.

Finally, we investigated the possibility that initially larger cardiomyocytes reach a size threshold faster and thus divide sooner than smaller cardiomyocytes. However, we observed no correlation between initial cardiomyocyte size and the timing of attempted cell division (**Fig. S2B-E**), regardless of ploidy. These results reveal that cardiomyocytes exhibit heterogeneous growth responses and may be able to modulate their individual growth rates in order to coordinate cell division within a defined time range post-stimulus. In our experiments, all cardiomyocytes analyzed attempted to divide within 30-36 hours on average in response to CHIR99021 treatment, regardless of ploidy. This is in line with a previous *ex vivo* imaging study estimating the total cell cycle length of ventricular cardiomyocytes to be around 30-35 hours (Hashimoto et al., 2014).

## DISCUSSION

As the advantages of digital holographic imaging are gradually being recognized, multiple applications of the Holomonitor M4 have recently been reported in various contexts such as wound healing, cell motility, three-dimensional culture of cells in matrigel, and optogenetics (Fang et al., 2023; Hellesvik et al., 2020; Tahara et al., 2018; Zhang & Judson, 2018). However, the application of digital holographic imaging in cardiovascular research has thus far been limited. The versatility of this instrument may have great potential to increase our cellular understanding of cardiomyocyte hypertrophy, migration, response to ischemia-reperfusion, and intercellular interactions. In this study, we first innovate a novel application of this technology to evaluate dynamic changes in cardiomyocyte size during hypertrophic growth and in response to a proliferative stimulus.

Current methods to directly evaluate cardiomyocyte size each have unique experimental limitations. For example, two-dimensional cardiomyocyte surface area measurements (Watkins et al., 2012) fail to consider changes in cell thickness and more directly evaluate cell spreading. Coulter counters (Li et al., 1996) and flow cytometry (Nakano et al., 2017) provide population-based measurements but do not resolve single-cell dynamics. And three-dimensional confocal imaging (Satoh et al., 1996) typically requires cell fixation and optical sectioning with high-powered lasers thus limiting long-term imaging of living cardiomyocytes.

We demonstrate that digital holographic imaging circumvents many of the limitations listed above and is a powerful methodology to quantitatively track primary cardiomyocyte size, ploidy, and proliferation dynamics in real-time with single-cell resolution. In rodents, mononucleated diploid cardiomyocytes attempt to divide, but fail cytokinesis, become binucleated, undergo cell cycle arrest, and increase in size within the first weeks of postnatal development (Bishop et al., 2021; Clubb & Bishop, 1984; Li et al., 1996; Soonpaa et al., 1996; Zak, 1973). The interplay between cardiomyocyte size, ploidy, and cell cycle regulation during this developmental transition is not well understood. Therefore, our unique application of digital holographic imaging technology is relevant to understanding the cellular mechanisms by which mammalian hearts lose their capacity for heart regeneration shortly after birth.

Our studies are the first to demonstrate that both mononucleated and binucleated cardiomyocytes increase in volume — 2.47-fold and 2.84-fold, respectively — prior to attempting cell division (**Fig. 5E**). Our findings suggest an intriguing possibility that neonatal cardiomyocytes control cell size and cell cycle progression based on a distinct variation of the “adder” model (Proulx-Giraldeau et al., 2022) where a constant amount of cell growth occurs during each cell cycle. Importantly, while the relative increase in size is conserved in both mononucleated and binucleated attempting cell division, binucleation increases the absolute volume of this threshold (**Fig. 5D**). This suggests that DNA content itself may directly regulate size dynamics in cycling cardiomyocytes. Cell growth has recently been reported to play an important role in cell cycle regulation by diluting the concentration of cell cycle inhibitors to trigger cell division (Zatulovskiy et al., 2020). It is possible that an increased copy number of genes encoding cell cycle inhibitors requires compensatory increases in cell volume in order to dilute inhibitor concentrations and permit cell division in binucleated cardiomyocytes.

Finally, we observed that while the majority of mononucleated cardiomyocytes successfully divided, more than half of binucleated cardiomyocytes did not attempt division in response to CHIR99021 stimulation (**Fig. 5B**). Importantly, these non-responsive cardiomyocytes exhibit a deficiency in growth during our 48-hour imaging experiments (**Fig. 5C**). These results support the conclusion that mononucleated cardiomyocytes retain high proliferative capacity and that binucleation limits the ability of cardiomyocytes to divide. This may occur by decreasing the ability of binucleated cells to reach a permissive size threshold required to attempt cell division. Interestingly, though binucleated cardiomyocytes attempted to divide with lower frequency, we did observe several cases of their successful division. Future work to better understand how the cell cycle is regulated in binucleated cardiomyocytes may lead to new strategies in cardiac regenerative medicine to trigger successful cell division in polyploid adult mammalian heart muscle cells.

While we demonstrate the powerful capabilities of the Holomonitor M4 digital holographic microscope in our study, we also note the technical limitations of the technology. For example, when measuring cellular optical volumes, the Holomonitor M4 and analysis software must identify true signal from background noise and there is some associated error in that measurement. This error is likely enhanced when evaluating smaller, thinner cells that tend to spread out on the cell culture surface (e.g., fibroblasts). Fortunately, primary cardiomyocytes tend to be larger, thicker cells (**Fig. 2B-C**), which lessens the impact of this error in our measurements. Another limitation is that the image segmentation and cell tracking features in the Holomonitor App Suite software does not perform well when cells are clustered very closely together. Therefore, accurate measurements using this approach relies on the identification of well-isolated single cells. This requirement would not be compatible with studies seeking to investigate cellular functions in the context of complex multicellular tissue environments. Nevertheless, our studies provide an example of how the strengths of the digital holographic microscopy can be leveraged to further understand the intrinsic cellular mechanisms that regulate cardiomyocyte proliferative potential.

## MATERIALS AND METHODS

### Animal breeding

Neonatal rat and mouse pups were generated through natural matings approved by Institutional Animal Care and Use Committee (IACUC) protocols 1072 (Dr. Payumo, San Jose State University) and AN197871-00B (Dr. Huang, University of California, San Francisco).

### Cardiomyocyte isolation and culture

Postnatal day 1 (P1) to P3 Sprague Dawley rat pups were euthanized by decapitation under manual restraint. Hearts were harvested and the aorta was pinched closed with forceps. The coronary vessels were then reverse-perfused with cardiomyocyte isolation buffer (CIB; 120 mM sodium chloride, 15 mM potassium chloride, 0.6 mM monopotassium phosphate, 0.6 mM sodium phosphate dibasic heptahydrate, 1.2 mM magnesium sulfate heptahydrate, 10 mM HEPES, 4.6 mM sodium bicarbonate, 30 mM taurine, 10 mM 2,3-butanedione monoxime, 5.5 mM glucose) supplemented with 0.4 mM EGTA by inserting a 20G needle into the left ventricle to clear out blood and improve digestion efficiency. After perfusion, the ventricles were isolated, cut into pieces, and transferred into a digestion buffer consisting of 2 mg/mL collagenase II (LS004176, Worthington Biochemical) and 0.3 mM calcium chloride in CIB. Tissues were digested at 37°C under gentle agitation for approximately 2 hours while collecting dissociated cells every 15 to 30 minutes until complete digestion. Collections were pooled and stored on ice to limit collagenase activity. After digestion, the dissociated cells were gently centrifuged for 5 minutes at 200 x g, resuspended in wash buffer 1 (CIB, 10% FBS, 0.64 mM calcium chloride), again in wash buffer 2 (CIB, 10% FBS, 1mM calcium chloride), and then in plating media composed of 10% FBS in DMEM/F12 (SH30023.01, Cytiva) with 100 µg/mL Primocin (ant-pm, InvivoGen). Cells were then passed through a 70-µm cell strainer (258368, Nest Scientific, NJ, USA) into a 6-well plate for pre-plating at 37°C in a 5% CO_2_ incubator for 1 to 2 hours to deplete non-cardiomyocyte populations via differential attachment. After pre-plating, the media and remaining cells were collected, quantified, and then seeded into 6-well plates (800,000 cells per well) or 12-well plates (400,000 cells per well) coated with 1% w/v gelatin (G9391, Sigma-Aldrich) in plating media overnight to allow cardiomyocyte attachment.

Neonatal mouse cardiomyocytes from ICR pups were isolated similarly with the following modifications. The digestion buffer additionally contained 0.2 mg/mL protease type XIV (P5147, Sigma-Aldrich), which may further help dissociate cell clusters into single cardiomyocytes. Also, 10% FBS was added prior to pooling each collection during digestion on ice to further protect the dissociated cells. Finally, after the preplating step, the cardiomyocyte-enriched cells were seeded into 96-well plates coated with 5 ug/cm^2^ rat tail collagen I (354236, Corning) in plating media at a density of 80,000-150,000 cells per well.

### Chemical treatments

Freshly isolated cardiac cells were cultured in plating media for 16-24 hours to allow attachment to the culture vessel. The next day, the cells were washed briefly three times in PBS to remove serum and then switched to a serum-free culture media composed of DMEM/F12 supplemented with 1X insulin, transferrin, and selenium (ITS, 400-145, GeminiBio) and 100 µg/mL primocin. At this time, cultures were additionally treated with 50 µM phenylephrine (P6126, Sigma-Alrdich) or 5 µM CHIR99021 (A133052, Ambeed) for 48 hours under serum-free conditions to stimulate cardiomyocyte hypertrophy or proliferation, respectively. When specified, cells were also incubated in 5 µM 5-ethynyl-2′-deoxyuridine (EdU, NE08701, Biosynth) for 48 hours to incorporate this thymidine analogue in any cell undergoing DNA synthesis within the treatment period.

### EdU detection

After chemical treatments, cardiac cultures were fixed in 10% formalin (15740-01, Electron Microscopy Sciences) for 15 minutes at room temperature. To detect EdU incorporation, cells were washed three times with PBS and blocked with 3% bovine serum albumin (A8412, Sigma-Aldrich) in PBS for 30 minutes at room temperature. Click chemistry was then performed to covalently attach an azide-containing fluorescent dye to the alkyne group available on EdU. In short, cells were treated with 10 µM FAM azide 6-isomer (A5130, Lumiprobe) in the presence of 1 mM CuSO_4_ and 250 µM ascorbic acid (AO537, Tokyo Chemical Industry) in 100 mM Tris pH 8.0 (AO321, Tokyo Chemical Industry) for 30 minutes.

### Immunohistochemistry

For rat cardiac cultures, samples were washed three times with 0.2% Triton X-100 in PBS (PBST) and then blocked in 10% FBS in PBST for 1 hour at room temperature directly after fixation or EdU detection. Cells were then incubated in cardiac troponin T antibody (MS-295-P1, Epredia) diluted 1:400 in PBST or alpha-actinin antibody (MA1-22863, Invitrogen) diluted 1:200 in PBST overnight at 4°C. The samples were again washed three times in PBST and then incubated in either Alexa Fluor 555 anti-mouse (A-31570, Thermo Fisher Scientific) or Alexa Fluor 488 anti-rabbit (711-545-152, Jackson ImmunoResearch) secondary antibodies diluted 1:500 in PBST for two hours at room temperature. Afterwards, the samples were washed three times in PBST with the first wash containing 1 μg/mL 4’, 6-diamidino-2-phenylindole (DAPI, AB2285492, Abcam) in order to stain nuclei prior to imaging on a Keyence BZ-X810 or Nikon TE300 inverted microscopes.

Mouse cardiac cultures were stained as above with the following modifications. Fixed cells were blocked in 5% normal donkey serum (NC9624464, Jackson ImmunoResearch) in PBST for 1 hour at room temperature prior to incubation in cardiac troponin T antibody. Nuclei were instead stained with 10 µg/mL Hoechst 33342 (5117, Tocris Bioscience) in the final washes of the protocol prior to imaging on a Nikon Eclipse Ti inverted microscope.

### Digital holographic time-lapse imaging

The Holomonitor M4 digital holographic microscope was purchased from Phase Holographic Imaging (PHI, Sweden) and placed directly within a Heracell 150 CO_2_ incubator (Thermo Scientific). For digital holographic imaging experiments, rat cardiac cells were cultured and treated with chemicals in 6-well plates (83.3920, Sarstedt) covered with HoloLids (PHI, Sweden) to stabilize imaging conditions. Live rat cardiac cultures were imaged on the Holomonitor M4 for 48 to 72 hours with time points taken every 20 min at 37°C and 5% CO_2_.

Mouse cardiac cells were cultured in a 96-well plate (83.3924, Sarstedt) for 2 days in plating media to allow attachment. Cells were then switched to a serum-free media composed of MCDB107 (E3000, United States Biological), 1X ITS (I3146, Sigma-Aldrich) and 100 µg/mL of Primocin. Prior to imaging on the Holomonitor M4, the plate was covered with HoloLids and images were acquired every hour for 5 days at 37°C and 5% CO_2_.

### Quantitative cell size and motility analyses

Time-lapse recordings were analyzed in the Holomonitor App Suite software (PHI, Sweden) using the “In-depth Analysis: Single Cell Tracking” feature to segment, trace, and quantify single cell dynamics. The following segmentation thresholds were applied to generate the results shown in Figures 2, 3, and Supplementary Figure 1: auto-minimum error method, adjustment 128, minimum object size 20. The following segmentation thresholds were applied to generate the results shown in Figures 4, 5, and Supplementary Figure 2: auto-minimum error method, adjustment 131, minimum object size 18. Cardiomyocytes were qualitatively distinguished from non-cardiomyocytes based on their larger size and limited motility as shown in Figure 2. The Holomonitor App Suite software was then used to track individual cells and optical volume, thickness, and motility measurements were taken at the specified time points. Care was taken to ensure that all cells analyzed were well-isolated and the accuracy of cell segmentation was manually verified on each imaging frame.

### Cardiomyocyte cell division analysis

72-hour time-lapse recordings were taken after stimulating the cardiac cultures to proliferate with CHIR99021. These recordings were analyzed in the Holomonitor App Suite software and well-isolated cardiomyocytes were identified and tracked using the “In-depth Analysis: Single Cell Tracking” feature. Mononucleated and binucleated cardiomyocytes were distinguished based on the number of visible nuclei identified within the cell. Within a 48-hour evaluation period, cardiomyocytes exhibiting complete furrow formation and separation into distinct daughter cells were considered to completely divide. Cardiomyocytes that attempted to separate but failed to resolve into two independent daughter cells were considered to incompletely divide. Cardiomyocytes that did not exhibit mitotic rounding during the 48-hour evaluation period were considered to have made no attempt at cell division. The extra 24 hours of the 72 hour time-lapse recordings was used to clarify cell division outcomes if necessary.

### Statistical analysis

All statistical analyses were performed using the GraphPad Prism software (GraphPad Software Inc.). Specific details including the statistical tests performed and the number of samples analyzed are reported in the corresponding figure legends.

## ACKNOWLEDGEMENTS

We thank Huang and Payumo laboratory members for their valuable advice on the manuscript. We thank Dr. Kersti Alm of Phase Holographic Imaging for technical advice. This paper was supported by American Heart Association Postdoctoral Fellowship (https://doi.org/10.58275/AHA.23POST1011296.pc.gr.161362) to S. Park, NIH awards (R01HL138456 and R01HL157280), American Heart Association Transformation Award and Established Investigator Award, and Tobacco-Related Disease Research Program Award P0558275 to Dr. G. N. Huang, and NIH SuRE-First award (R16GM146643) to Dr. A. Y. Payumo.

## AUTHOR CONTRIBUTIONS

SP, HH, GNH, and AYP designed experiments. SP, HH, and IR, performed all experiments. SP, HH, IR, GNH., and AYP analyzed the data. JM, SK, and JS helped to optimize experimental conditions. SP, HH, GNH, and AYP drafted the manuscript. SP, HH, IR, JM, SK, JS, GNH, AYP edited the manuscript.

## SUPPLEMENTARY MATERIALS

**Figure S1:**
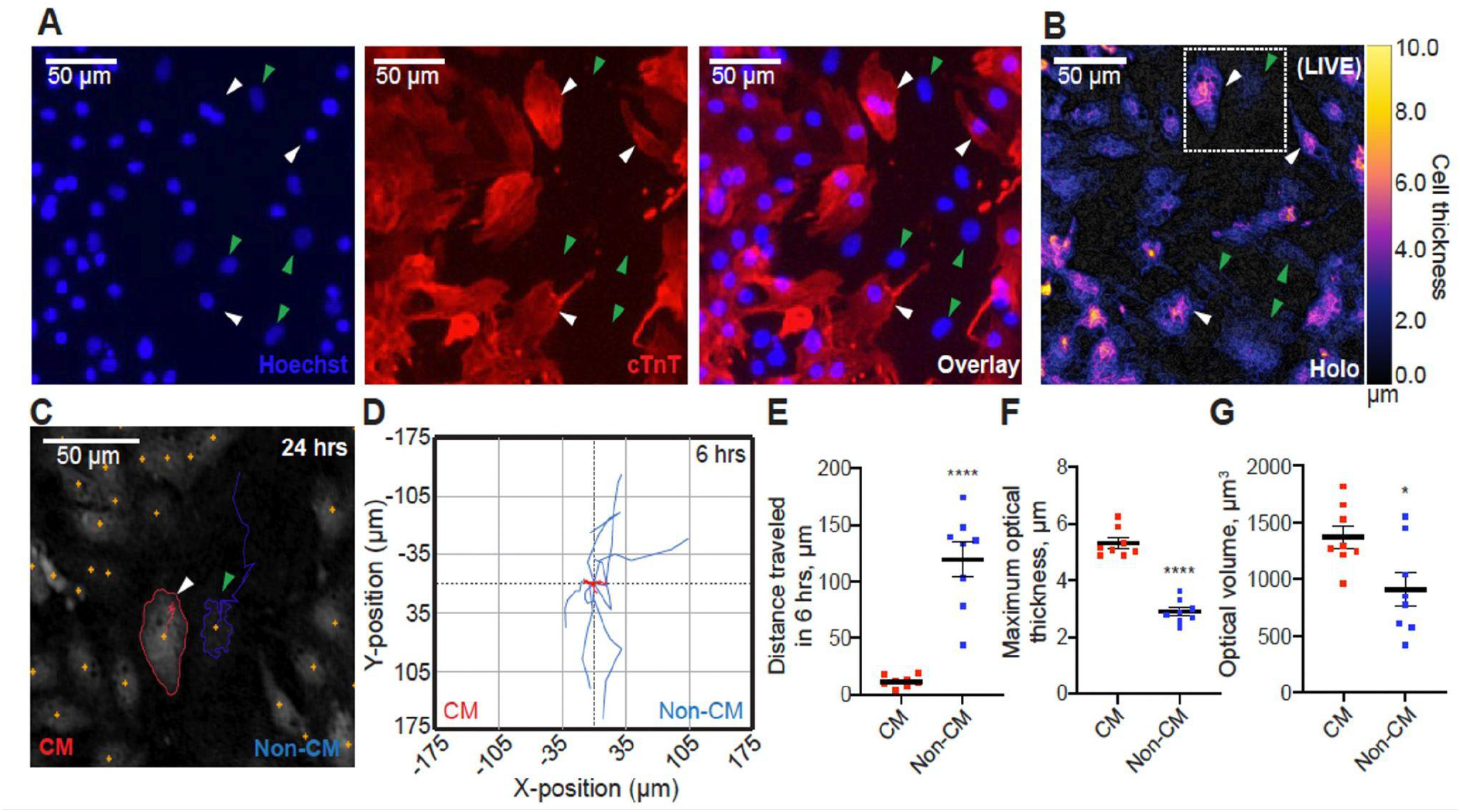
Label-free digital holographic imaging can also identify primary mouse cardiomyocytes based on their larger size and limited motility. (**A-B**) The same imaging field of fixed mouse cardiac cells visualized on a phase contrast fluorescence microscope (**A**) or live on the Holomonitor M4 (**B**). Cardiomyocytes (CM) are cardiac troponin T (cTnT)-positive (**A**, white arrowheads) and larger (**B**, white arrowheads) while non-CMs are cTnT-negative (**A**, green arrowheads) and smaller (**B**, green arrowheads). Hoechst labels all nuclei. (**C**-**D**) Motility analysis of CMs and non-CMs. (**C**) Representative segmentation and tracking of a CM (red outline, white arrowhead) and non-CM (blue outline, green arrowhead) shown in the dotted square in **B** over 24 hours. Cell trajectories are shown as a single solid line for each cell. (**D**) Overlaid trajectories of CMs (n=8) and non-CMs (n=8) over 6 hours. All cell traces begin at the (0,0) coordinate. (**E**) Total distance traveled by each cell tracked in **D** over 6 hours. (**F-G**) Maximum optical thicknesses (**F**) and optical volumes (**G**) of the same cells tracked in **D**. Values are reported as mean±SEM. Student’s t-test; *, *P* < 0.05; ****, *P* < 0.0001.

**Figure S2:**
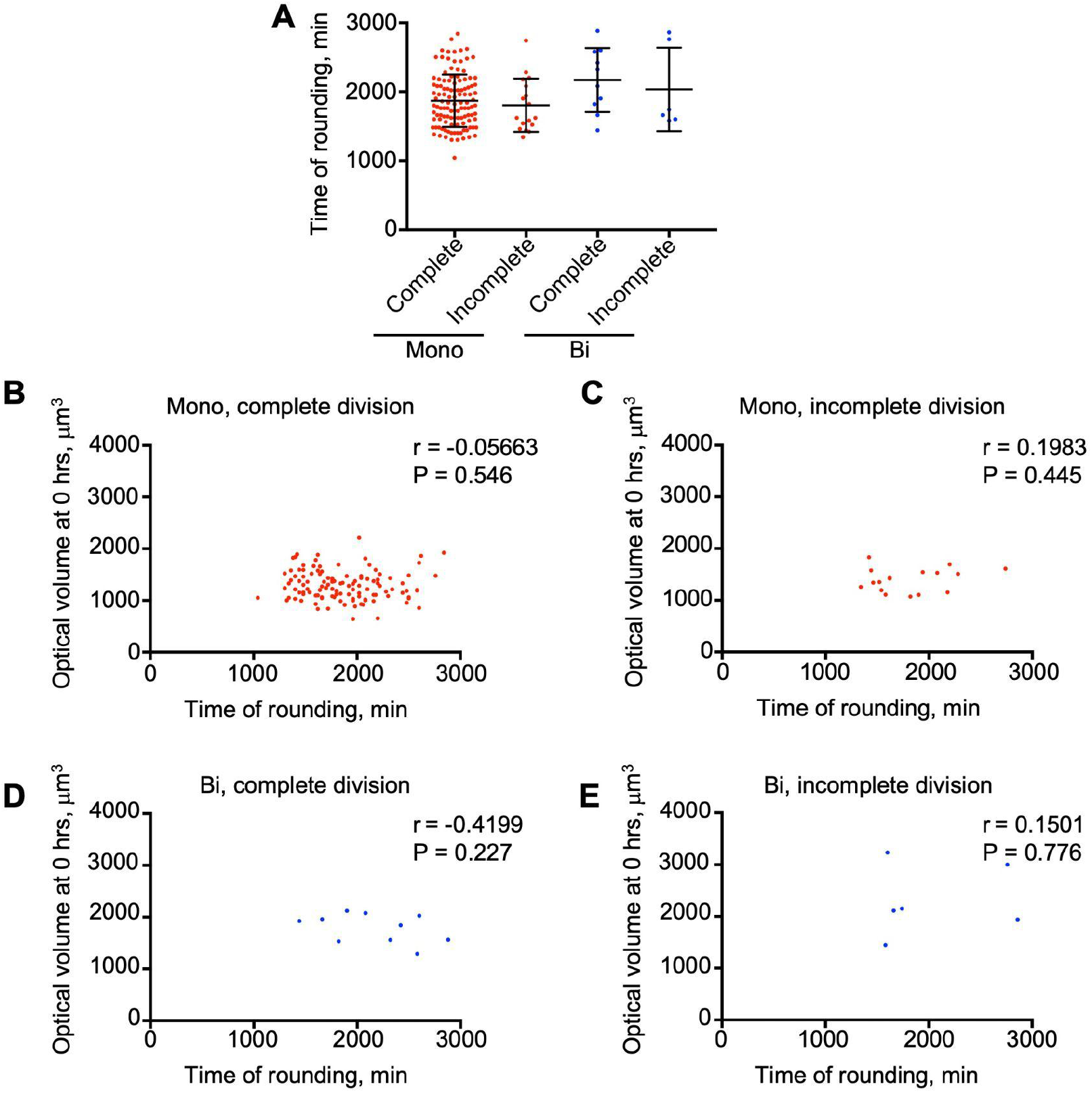
Initial cardiomyocyte size does not correlate with the timing of attempted cell division. (**A**) Time of rounding for postnatal day 3 CMs that attempt to divide after stimulation with CHIR99021 (mononucleated, complete, n = 116; mononucleated, incomplete, n = 17; binucleated, complete, n = 10; binucleated, incomplete, n = 6). One-way ANOVA revealed no significant differences. (**B-D**) Correlation plots of initial CM size and time of rounding. Pearson’s correlation coefficient, r, and P-values, P, are shown demonstrating no significant correlations.

